# The constitutive oncogenic and signaling activities of phosphatidylinositol 3-kinase (PI3K) isoforms p110β and p110δ

**DOI:** 10.1101/2025.06.26.659600

**Authors:** Yoshihiro Ito, Petra Pavlickova, Claire S. Levy, Sohye Kang, Jonathan R. Hart, Lynn Ueno, Peter K. Vogt

**Affiliations:** The Scripps Research Institute, Department of Molecular and Cellular Biology, 10550 N. Torrey Pines Road, La Jolla, CA 92037, USA; private practice; Bristol Myers Squibb, San Diego, CA; University of California, San Francisco, CA; TAE Life Sciences, Santa Monica, CA; U.S. Department of State, Washington, D.C

**Keywords:** domain analysis, oncogenic activity, signaling activity, class IA PI3K, isoforms

## Abstract

The p110β and p110δ isoforms of the catalytic subunit of phosphatidylinositol 3-kinase (PI3K) show enhanced oncogenic and signaling activities as compared with the p110α protein. The adapter binding domains (ABDs) of p110β and p110δ contain an isoform-specific PXXP motif. Mutations of this PXXP to AXXA diminish the oncogenic and signaling activities. This loss of function can be compensated by placing a gain-of-function mutation in the helical domain of the P/A mutants. The P/A mutants still associate with the regulatory p85 subunit, but the affinity of this interaction is decreased. p110α with the ABD of either p110β or p110δ shows an increase of oncogenic and signaling activities, whereas p110β or p110δ with the ABD of p110α have greatly reduced oncogenic and signaling activities. Introducing the PXXP motif in the ABD of p110α results in a significant gain of function. We conclude that the PXXP motif in the ABD of p110β and p110δ is essential for the elevated oncogenic and signaling activities of these isoforms. We propose that the PXXP motif in the ABD of p110β and p110δ affects the interaction with the iSH2 domain of p85, shifting the regulatory SH2 domains into less effective inhibitory conformations.

## 1. Introduction

Phosphatidylinositol 3-kinases (PI3Ks) comprise a family of dimeric enzymes that in higher eukaryotes is divided into three classes [1–3]. Class I A of PI3K encompasses three isoforms of the catalytic subunit, p110α, p110β and p110δ [4]. These isoforms play distinct roles in cellular signaling and in the regulation of cell growth [5–14]. In cancer, p110α shows consistent somatic gain-of-function mutations [15], and these mutations are oncogenic in cell culture and in animals and act as cancer drivers [16, 17]. The isoforms p110β and p110δ are much less frequently mutated in human tumors but can show differential expression and other links to oncogenicity [18–32]. In the area of PI3K inhibitors, there has been a growing effort toward isoform specificity [5, 33–39]. The currently FDA-approved PI3K inhibitors show various degrees of this specificity [40–44].

One of the puzzling isoform-specific properties of PI3K is the enhanced signaling and oncogenic activity of the wild type proteins p110β and p110δ. They readily induce cellular transformation accompanied by high PI3K signaling activity, whereas overexpression of wild type p110α is not oncogenic in cell culture [45]. This phenomenon has been analyzed by structural and by genetic approaches [46, 47]. In the present study, we focus on sequence hallmarks shared by p110β and p110δ and identify an isoform-specific PXXP motif in the adapter binding domains (ABDs) of p110β and p110δ. In p110α, the PXXP motif is absent. We mutated the prolines in this motif to alanines, generating p110β P104A/P107A and p110δ P94A/P97A. These mutations will be referred to collectively as P/A mutations or P/A mutants. Our study shows that the PXXP motif is required for the unique oncogenic potential of p110β and p110δ. The enhanced oncogenic transforming activity can be transferred to p110α by exchanging its ABD for that of p110β or p110δ or, to a lesser degree, introduction of the PXXP motif. We hypothesize that the PXXP motif controls a conformation that favors constitutive activity of the protein.

The class I A isoforms of PI3K, their domain structure and the mutations used in this study are depicted in Fig. 1.

**Figure 1.**
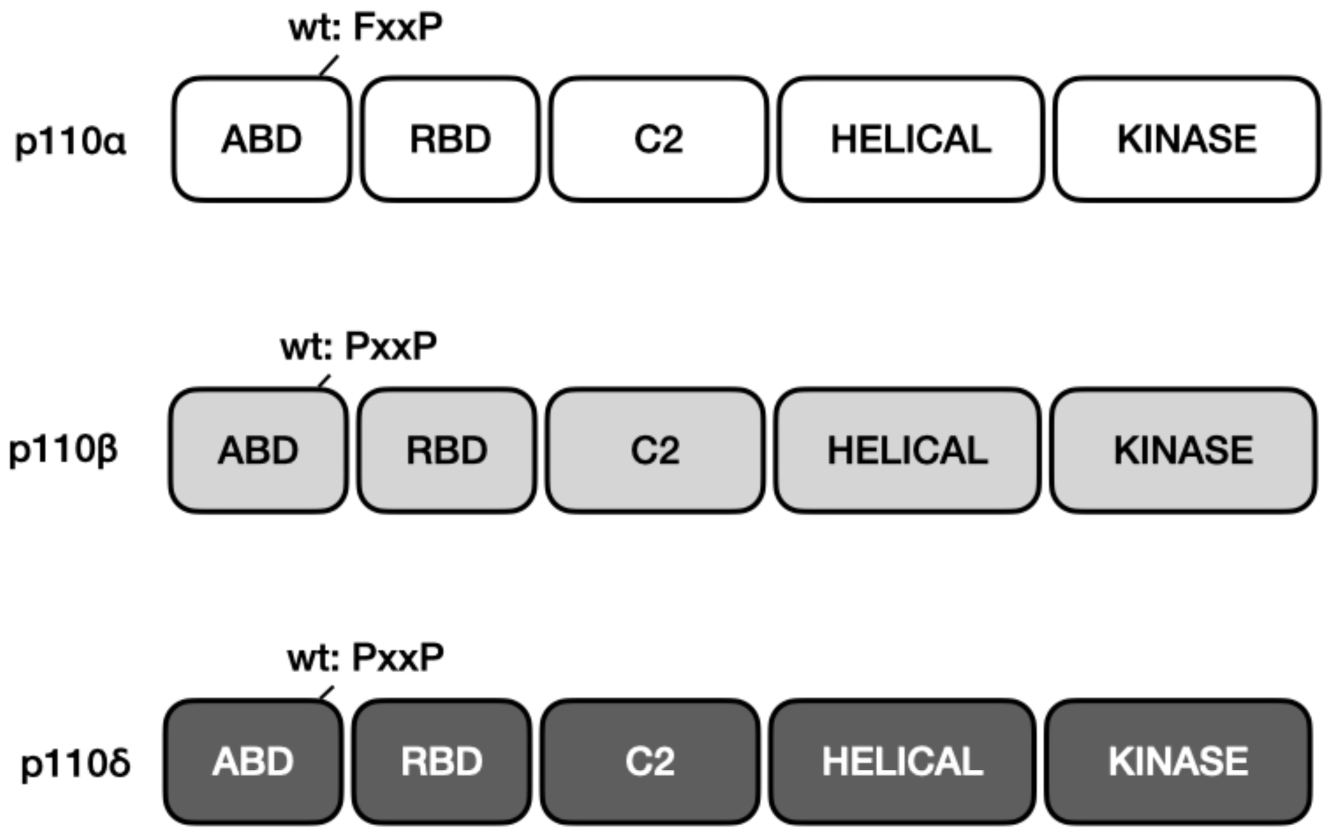
Class I A PI3K catalytic subunits. Schematic representation of domain structure from N to C terminus. ABD = adapter binding domain, RBD = Ras binding domain, C2 = C2 domain, helical and kinase domains. The location of the PXXP motif in p110β is residue 104 – 107; in p110δ it is residue 94 – 97.

## 2. Results

### 2.1. The P/A mutants of p110β and p110δ have lost the enhanced oncogenic and signaling activity of the wild type protein but the oncogenic phenotype can be rescued by a gain-of-function mutant in the helical domain

As a first test of the significance of the PXXP motif, we compared oncogenic and signaling activities of the P/A mutants with their wild type progenitors. The P/A mutants have lost most, but not all of the cell-transforming and AKT-phosphorylating activity of the wild type proteins (Fig. 2 A, B). This loss can be compensated by specific mutations in the helical domains of p110β [E552K] and p110δ [E525K]. These are analogous to common hotspot gain-of-function mutations in p110α that eliminate an electrostatic interaction between p110 and the N-terminal SH2 domain of p85 of PI3K. They also have an activity-enhancing effect in p110β [26] and p110δ, and a similar mutation of p110β has been identified in breast cancer [27]. Adding the helical domain mutation to the P/A mutation leads to a restoration of oncogenic and signaling activities in the combined mutants (Fig. 2 A, B). These data suggest that the PXXP motif is required for the enhanced oncogenic and signaling activities of wild type p110β and p110δ, and the effects of its mutational loss can be largely compensated by an additional mutation in the helical domain.

**Figure 2.**
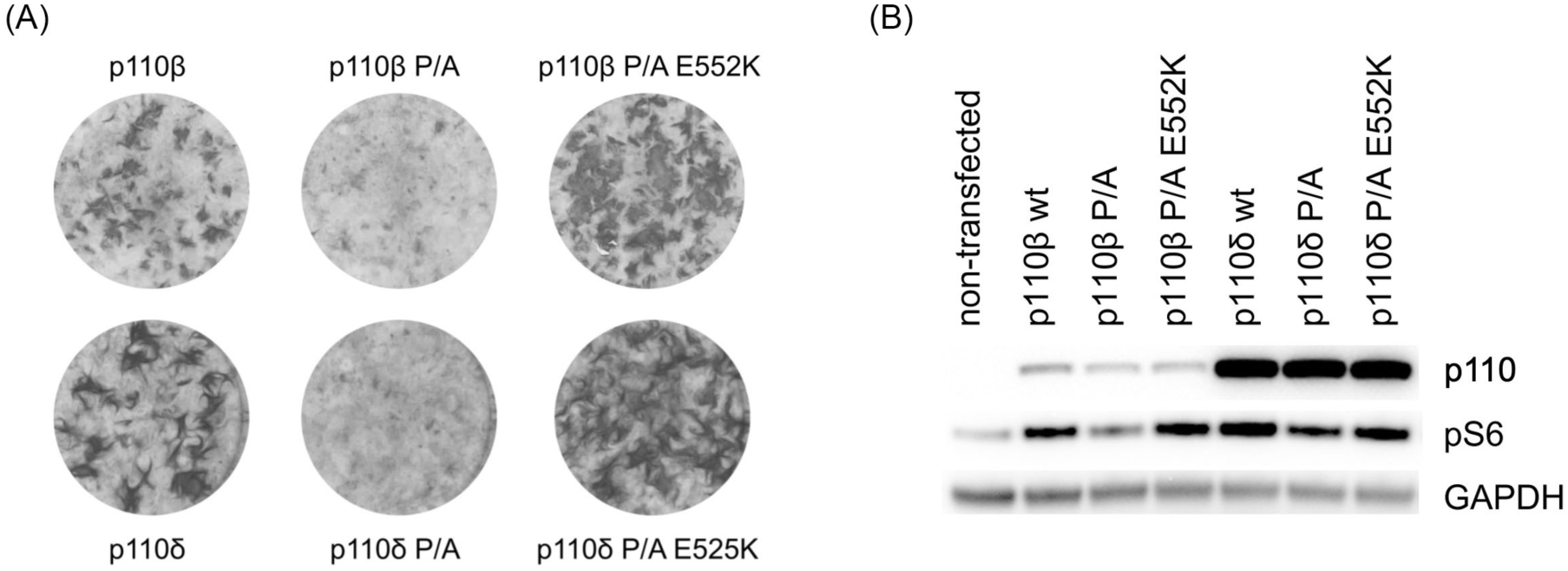
(A) Oncogenic transformation of CEF by wild type p110β and p110δ and their mutants. Transformation activity is strongly reduced by the P/A mutations in the ABD and is rescued by the E/K mutations in the helical domain. The photographs represent cell cultures transfected with 0.5 µg/well of the RCAS(A) vector expressing the p110 proteins. (B) Display of PI3K signaling by Western blot using phosphorylation of the ribosomal protein S6 as indicator. (In this Western blot, untagged p110β was used and was detected by isoform-specific antibody.)

### 2.2. The P/A mutants show reduced binding to p85, and this interaction is virtually abolished by the addition of the helical domain mutation

In order to detect interactions between wild type p110, its mutants and p85, we N-terminally FLAG-tagged p110α, p110β and p110δ and C-terminally HA-tagged p85α and p85β. The proteins were expressed from a lentiviral vector in 293T cells as described in the Materials and Methods section. The associations between the p110 and p85 proteins were visualized by co-immunoprecipitation (Fig. 3). For p85β, binding to the P/A mutants is reduced and becomes negligible for the combined P/A helical domain mutants. For p85α, the binding data to the P/A and P/A plus helical domain mutants are less compelling but point in the same direction.

**Figure 3.**
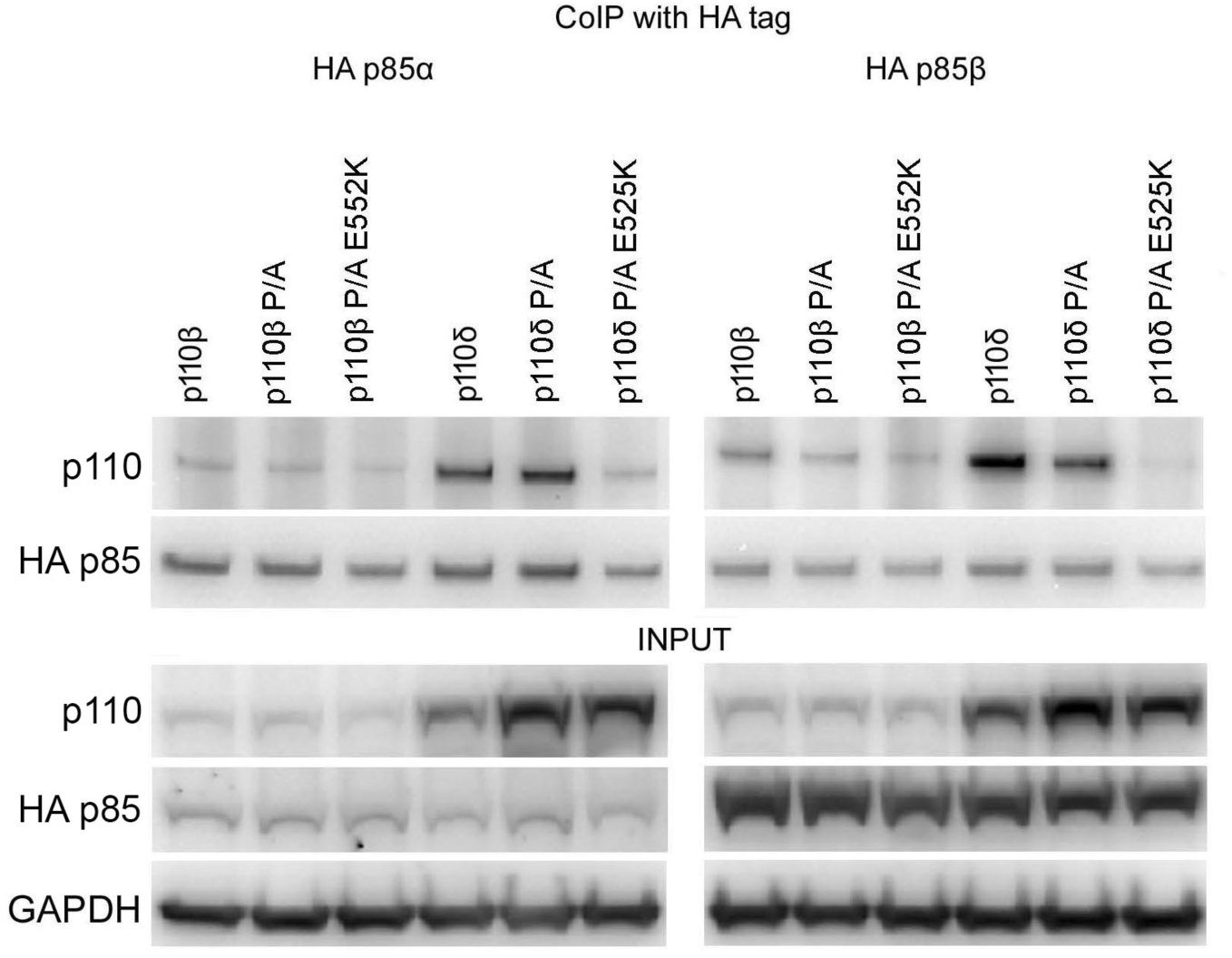
Interaction of the P/A mutants of p110β and p110δ and of the P/A plus helical mutants with p85α and p85β. p85 was HA-tagged, and p110 was FLAG-tagged; cell lysates were immunoprecipitated with anti-HA-tag antibody. FLAG-tagged p110 was visualized with anti- FLAG antibody by immunoblotting.

### 2.3. Deletion of the ABD increases p110 expression and activity

As a complementary experiment to the mutations in the ABD and the helical domains, we deleted the ABD from the three isoforms of class I A PI3K and determined the effect of these deletions on expression and on oncogenic and signaling activities (Fig. 4). In this experiment, the wild type and the truncated proteins were expressed in chicken embryo fibroblasts (CEF) by the RCAS(A) retroviral vector as described in the Materials and Methods section. For all three isoforms, the expression levels of the truncated proteins showed a significant increase compared to the levels of the wild type progenitors. This finding is in accord with previous observations [26, 48]. The lack of interaction between the ABD-deleted p110 isoforms and p85α as well as p85β was confirmed by co-immunoprecipitation. In these experiments, the C-terminally FLAG-tagged p110 proteins and the HA-tagged p85α and p85β were expressed in 293T cells from a lentiviral vector, and co-immunoprecipitation was performed as detailed in the Materials and Methods section. The increase of expression seen with the ABD-deleted p110 isoforms was correlated with enhanced oncogenic transformation of CEF and with slightly but significantly increased signaling as indicated by the level of phosphorylated ribosomal protein S6. Our data do not permit a distinction between enhanced protein-intrinsic activities and increases of recorded activity that reflect elevated pro- tein levels.

**Figure 4.**
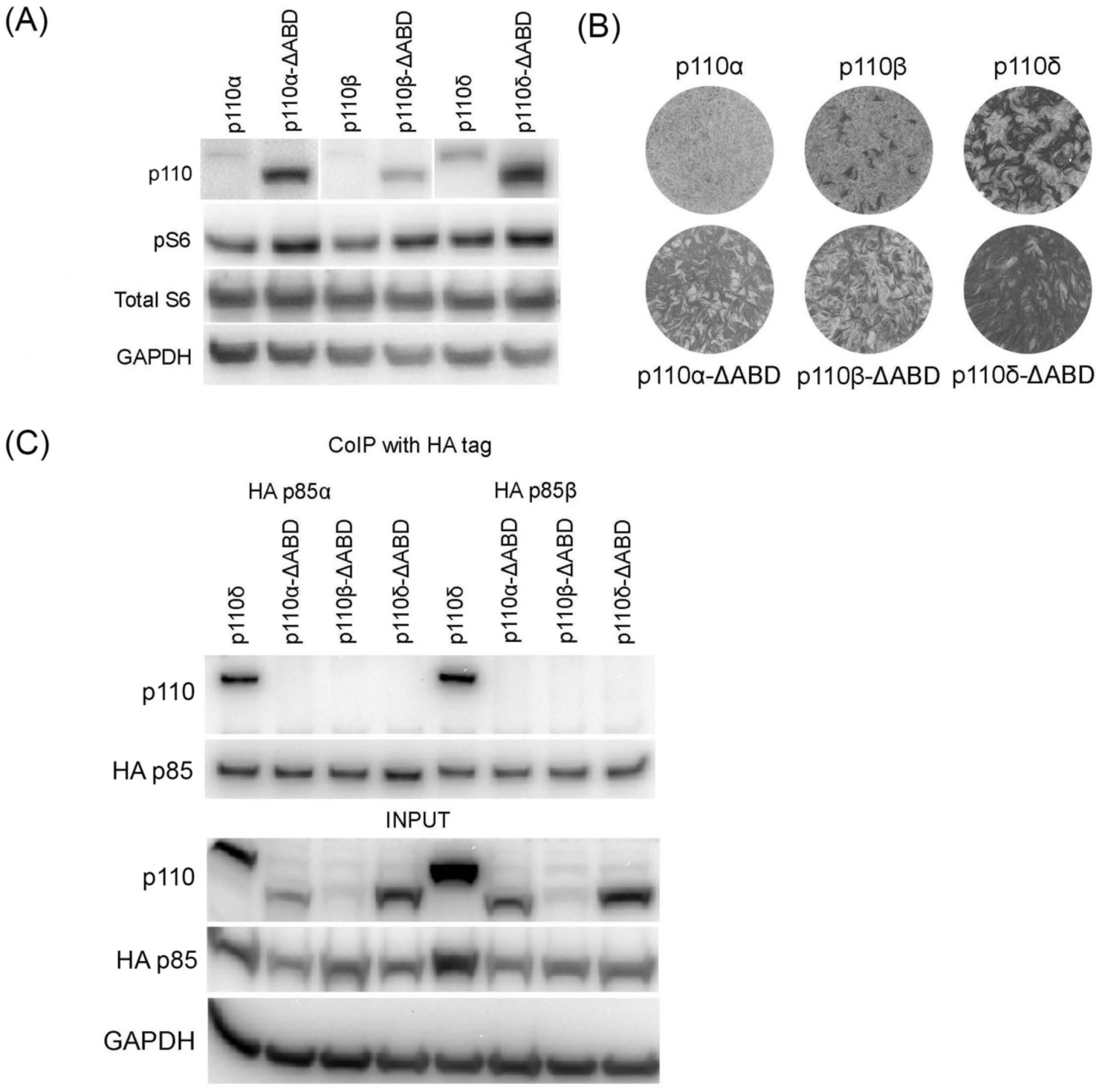
(A) The ABD-deleted FLAG-tagged p110 proteins showed increased levels of expression compared to wild type controls as visualized by immunoblotting with anti-FLAG antibody. The increased expression is correlated with increased PI3K signaling represented by phosphorylation of S6. (B) Truncation of the ABD enhances oncogenic activity in p110α, p110β, and p110δ. The photographs represent cell cultures transfected with 0.5 µg/well of the RCAS(A) vector expressing the respective p110 proteins. (C) The ABD-truncated p110α, p110β and p110δ do not bind p85α or p85β.

### 2.4. The P/A-mutated ABD of p110β shows extremely low binding to p85

We also investigated the ability of the isolated ABDs to interact with p85 by co-immunoprecipitation (Fig. 5). The ABDs of p110α and p110β were C-terminally FLAG- tagged and expressed with HA-tagged p85 via a lentiviral vector in 293T cells. The analysis followed the general scheme used for Figs. 3 and 4 and is detailed in the Materials and Methods section. The wild type ABDs showed definitive interaction with p85α. The interaction with p85β was strong for the p110α ABD but was reduced for the p110β ABD. The P/A-mutated ABD of p110β showed very low (p85α) or undetectable binding (p85β). These observations are in accord with a reduced interaction of P/A-mutated p110β with both p85 isoforms as suggested in Fig. 3. Much of the residual binding between P/A-mutated p110 and p85 seen in the experiment depicted in Fig. 3 appears to be mediated by contacts outside the ABD.

**Figure 5.**
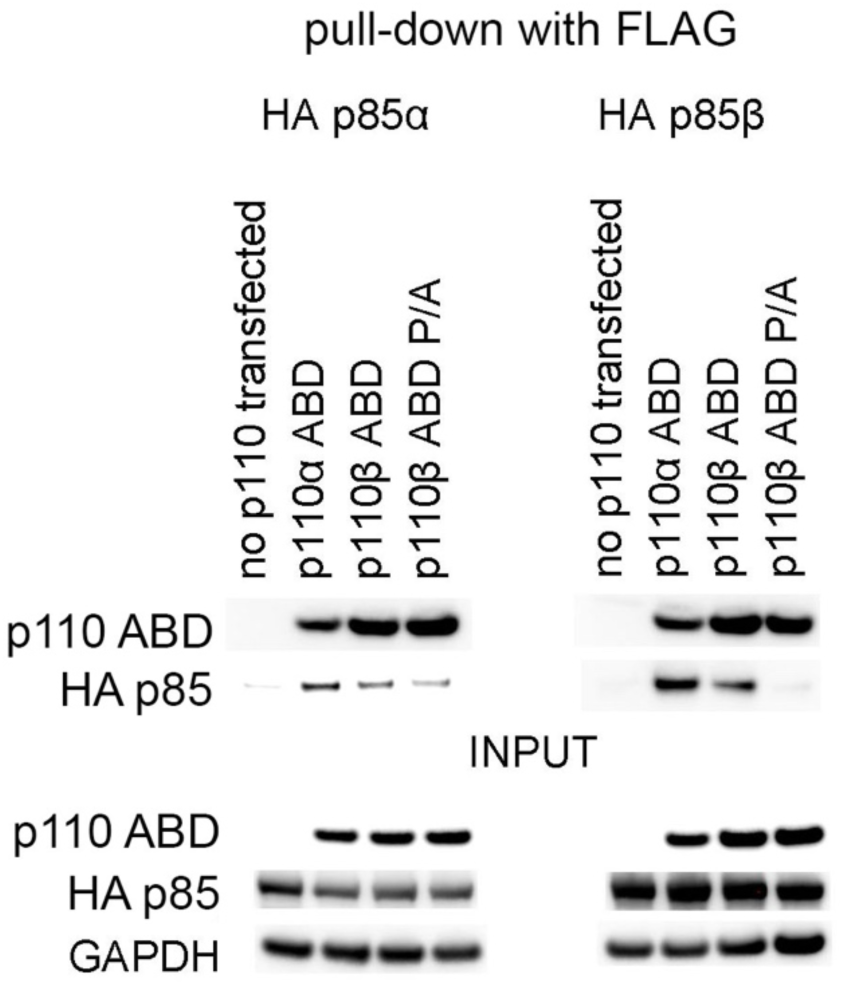
The isolated ABD of p110α and of p110β still bind to p85α and p85β, but the p110β ABD carrying the P/A mutations shows only very weak or no binding to p85. Immunoprecipitation and immunoblotting were performed as described in Figure 3.

### 2.5. Enhanced oncogenic and signaling activity can be transferred to p110α with the p110β or p110δ ABD

We further investigated the role of the PXXP motif in enhancing the oncogenic activity of wild type p110β and p110δ by exchanging the wild type and P/A-mutated ABDs among the p110 isoforms of class I A PI3K (Fig. 6). The p110α constructs carrying the p110β or p110δ ABDs show increased oncogenic activity, comparable to that of wild type p110β and p110δ. They also mediate more active signaling as indicated by the level of phosphorylated S6. In contrast, p110β and p110δ with the α ABD show loss of oncogenic and signaling activity. The P/A-mutated ABDs of p110β and p110δ also enhance oncogenic activity, but to a significantly lesser degree than the wild type versions. This activity is in accord with the observation that the P/A mutations reduce but do not extinguish the activities of the wild type proteins.

**Figure 6.**
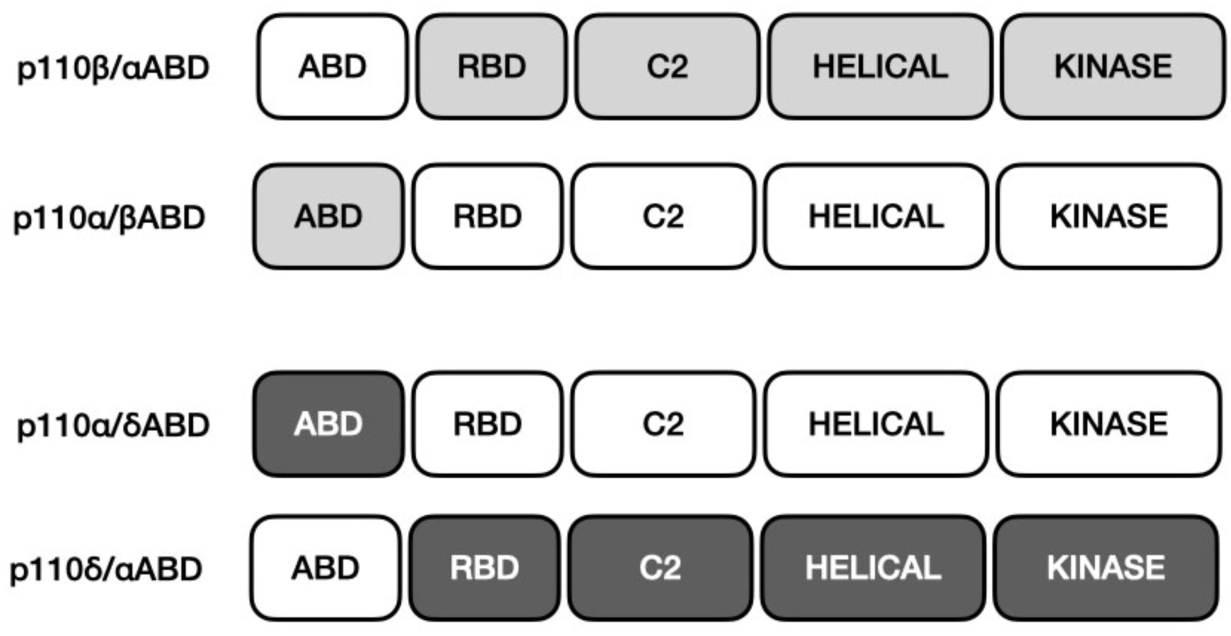
Chimeras of p110 generated by reciprocal exchanges of the ABDs of p110α, p110β, and p110δ.

It could be argued that the chimeric constructs between p110α and the ABDs of p110β and p110δ are conformationally altered, so that they no longer interact with p85. The enhanced activities could then reflect the same type of activation that is seen with the ΔABD constructs (Fig. 4). In this situation, enhancement of activity would not be an intrinsic property of the β and δ ABDs. We tested this possibility by co-immunoprecipitation (Fig. 7). All ABD chimeras bind p85, supporting the conclusion that their positive and negative effects on oncogenic and signaling activities are domain-intrinsic properties.

**Figure 7.**
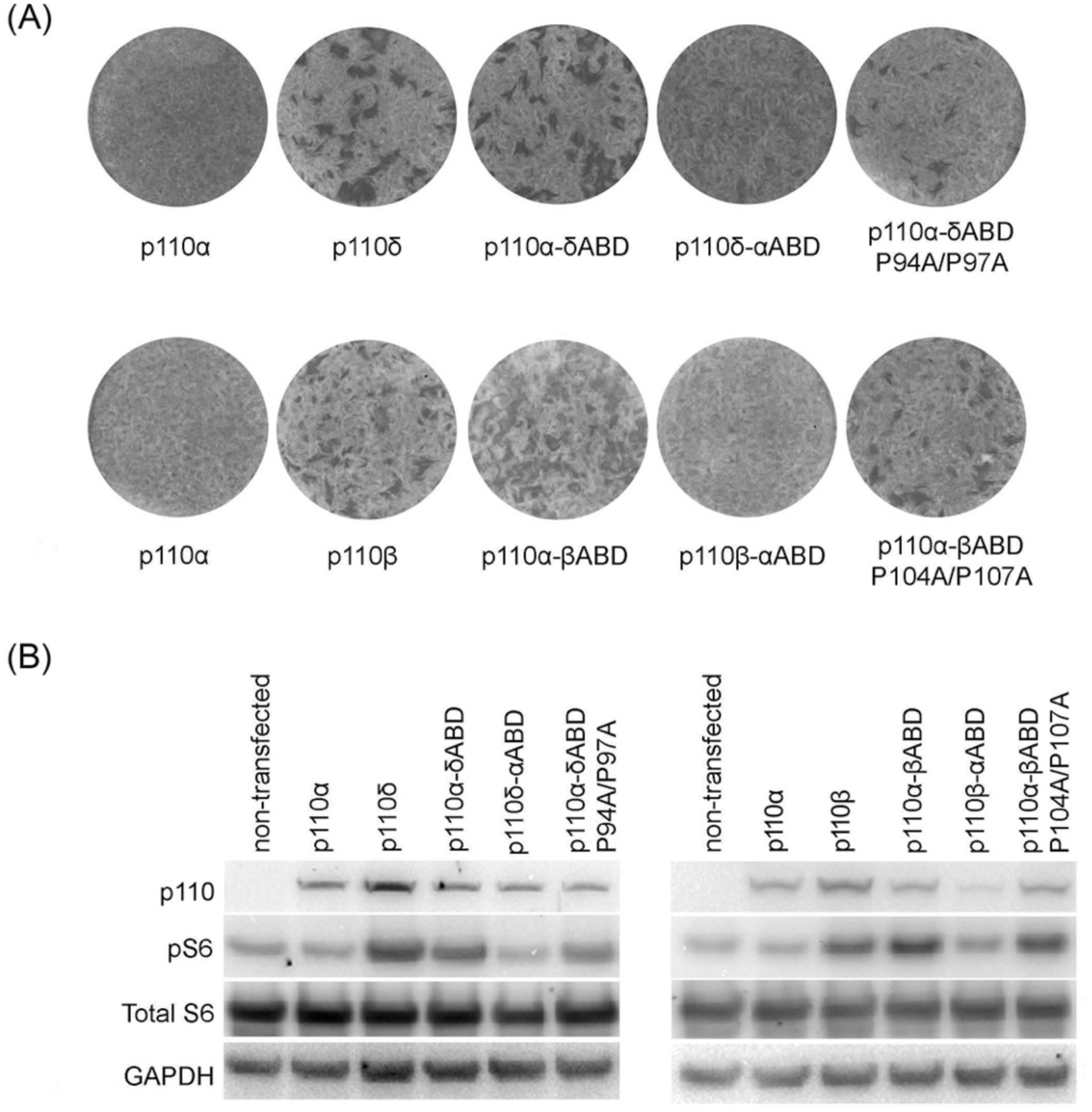
Reciprocal exchange of ABDs between p110α and p110δ and between p110α and p110β. (A) The photographs show oncogenic transformation in cell culture induced with 0.5 µg DNA of the RCAS(A) vector expressing the respective proteins. The ABDs of p110β and p110δ confer strong transforming activity on p110α, and the ABD of p110α does the reverse, greatly reducing oncogenic activity of p110β and of p110δ. The ABD of p110β carrying the P/A mutation and the ABD of p110δ carrying the P/A mutation moderately enhance the oncogenic activity of p110α. (B) The effect of the domain exchanges on oncogenic activity also extends to signaling as indicated by the phosphorylation of S6.

### 2.6. The oncogenic and signaling activities of p110β and p110δ are not correlated with protein expression levels or stability

Multiple factors affect levels of protein expression mediated by the RCAS(A) or lentiviral vectors. One of these is protein stability which in turn could affect the measured activity of the protein. We investigated this possibility for four constructs of p110δ: the wild type, the ABD deletion mutant, the P/A mutant and the combined P/A-helical do main mutants. The C-terminally FLAG-tagged proteins were expressed in CEF with the RCAS(A) retroviral vector and analyzed by immunoblotting using a FLAG tag antibody. Protein stability was determined by cycloheximide treatment as described in Materials and Methods (Fig. 8). Wild type p110δ, p110δ-ΔABD as well as the combined P/A-helical domain mutants are highly active in oncogenic transformation and signaling. The P/A mutant shows greatly reduced activity. These activities are not correlated with protein stability. The findings are in agreement with observations on protein levels made in the course of this study. For instance, p110β and its mutants are generally expressed at a lower level than p110δ (Figs. 3, 4, 7). Yet, these differences are not reflected in oncogenic or signaling activity. We conclude that the oncogenic and signaling activities determined in the current investigation do not simply reflect protein levels or protein stability. The data in Fig. 9 are also in agreement with the p110-stabilizing activity of p85. The p110 constructs that fail to interact with p85 are significantly less stable than the interacting p110 constructs. Yet, despite its reduced stability, the ABD-truncated p110δ shows no decrease in oncogenic or enzymatic activity (Fig. 4).

**Figure 8.**
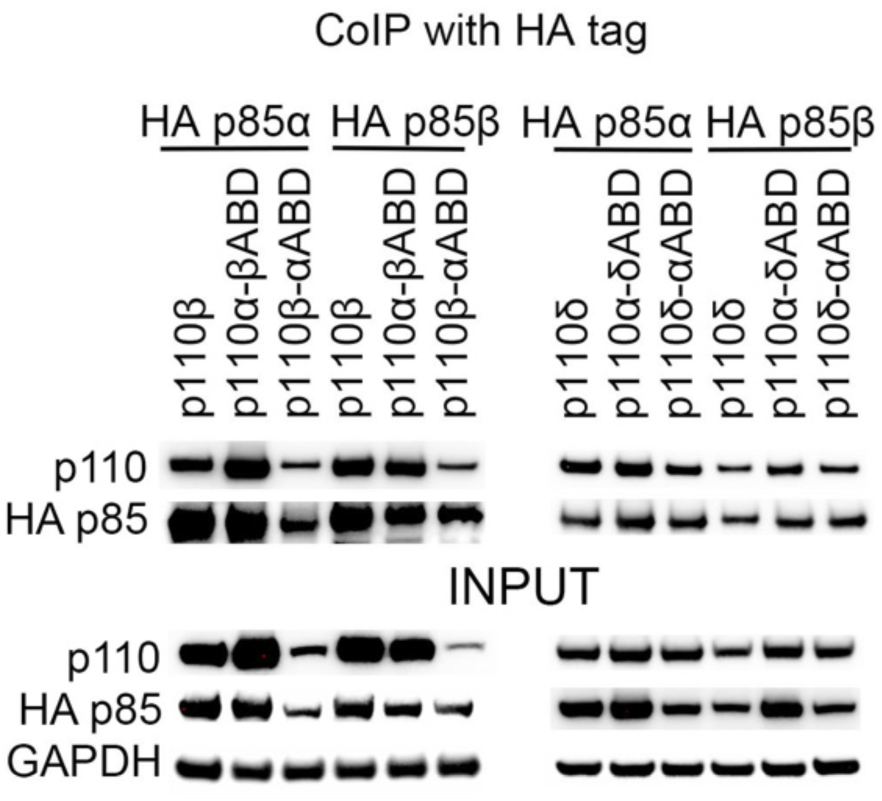
Co-immunoprecipitation with HA-tagged p85. All ABD chimeras still bind to p85.

**Figure 9.**
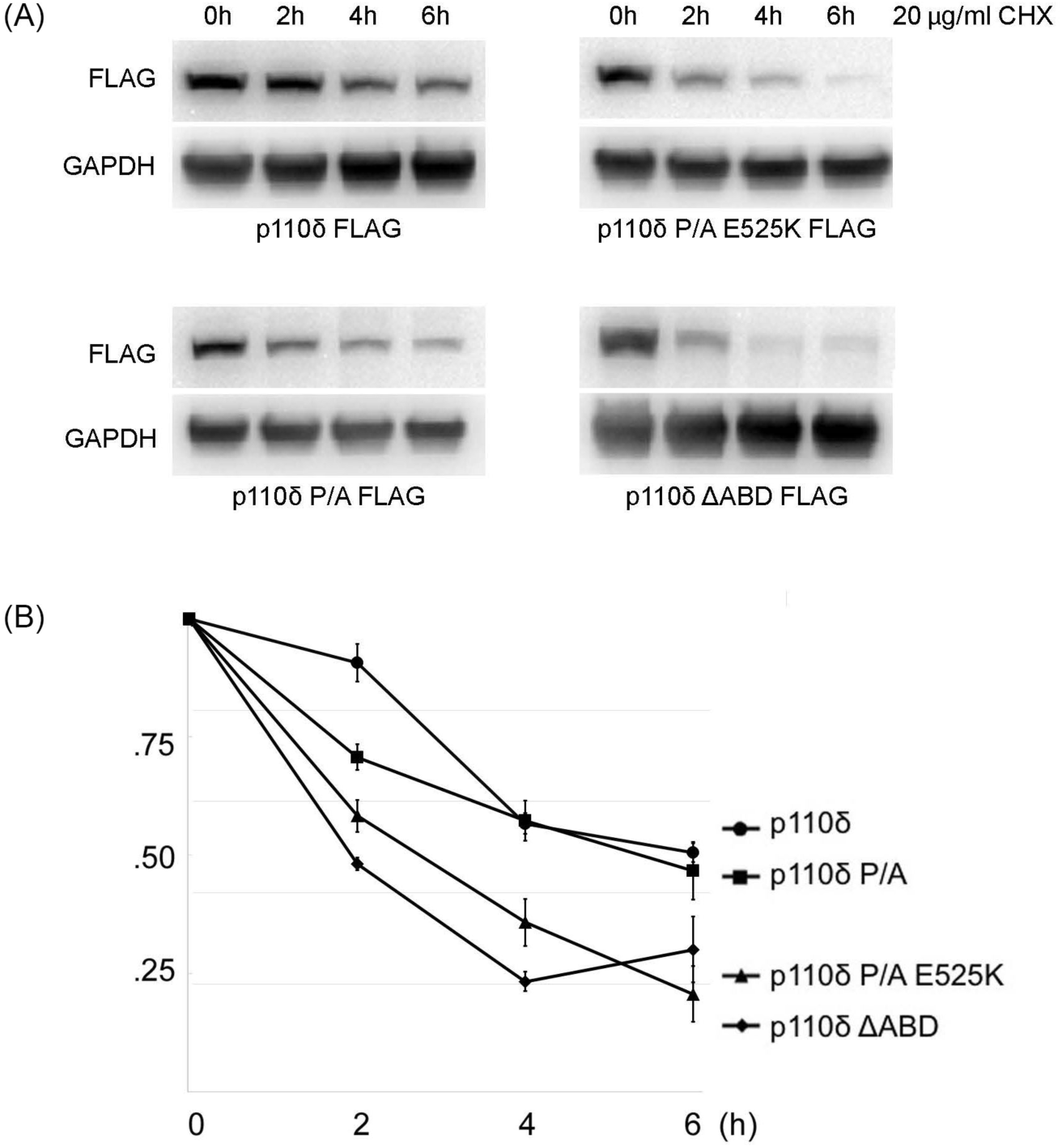
Oncogenic and signaling activities are not correlated with protein stability. (A) Cells expressing p110δ, p110δ P/A, p110δ P/A E525K and p110δ ΔABD were incubated with cycloheximide (CHX, 20 mg/ml) for the indicated times. Protein levels were analyzed by immunoblotting. (B) Time course of p110 protein degradation.

### 2.7. Introduction of a PXXP motif into the ABD of p110α induces a gain of function

We mutated wild type p110α to generate a PXXP motif at a position that is presumably analogous to that in the ABD of p110β and p110δ. Specifically, the residues 94 to 97 in wild type p110α are LFQP, and these were changed to PFLP (L94P, Q96L). The mutant was expressed in CEF for an estimate of oncogenic and signaling activities. Fig. 10 shows a small, but significant increase in signaling as measured by the phosphorylation of S6 in mutant-expressing cells compared to cells expressing wild type p110α. The mutant also acquired the ability to induce oncogenic transformation as evidenced by focus assay in CEF. Again, this gain of function was small compared to the activities of p110β and p110δ, but it was significant. We conclude that the introduction of a PXXP motif in the ABD of p110α induces gains of function in signaling and oncogenic activity.

**Figure 10.**
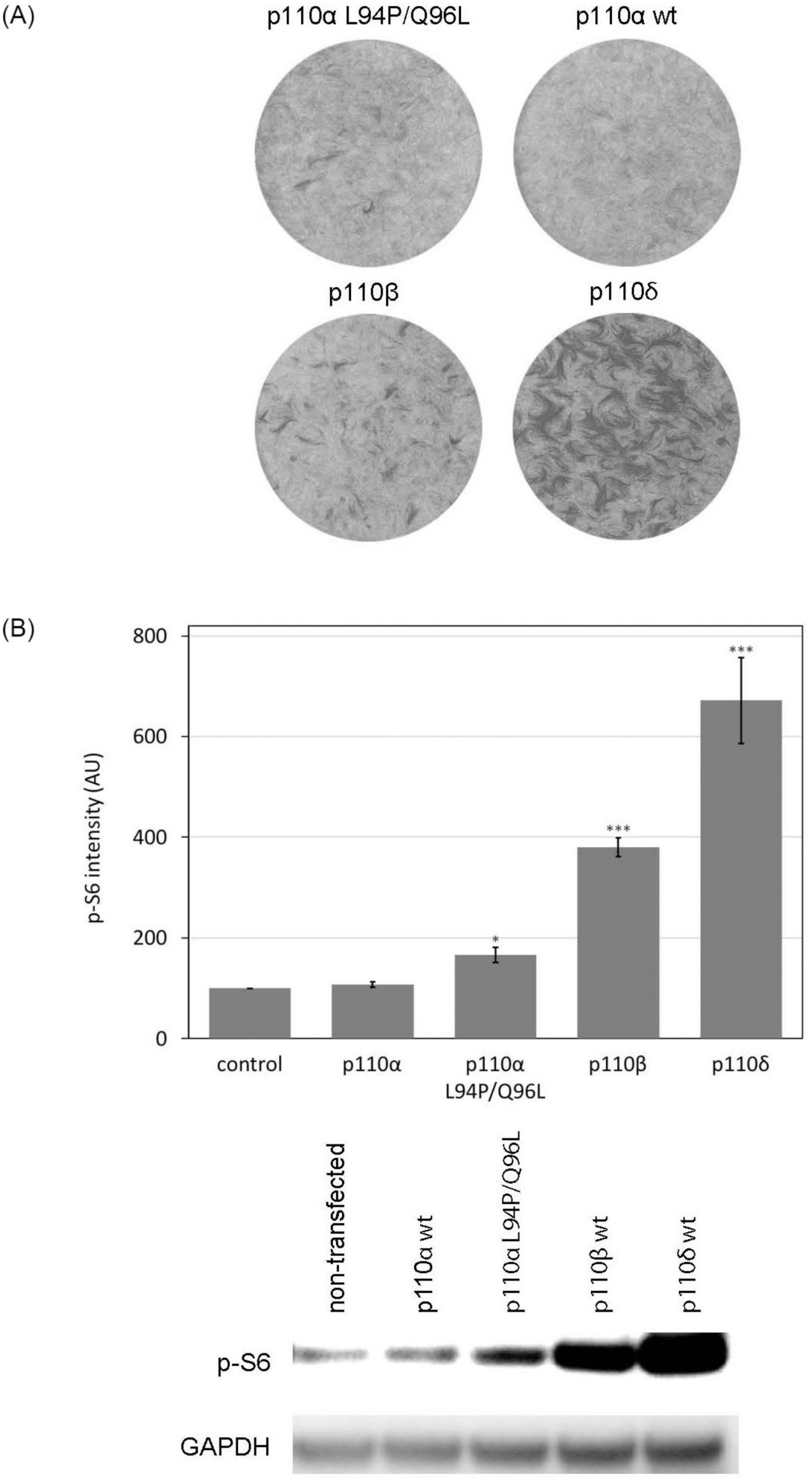
Introduction of the PXXP motif in the ABD of p110α induces a gain of function. (A) Oncogenic activity as measured by transformation of CEF. Representative photographs of cell cultures transfected with 0.5 µg/well of the RCAS(A) vector expressing the respective p110 proteins. (B) Integrated band intensities of p-S6 staining of CEFs that express the indicated RCAS constructs. * p < 0.05, ** p <0.01, *** p < 0.005

## 3. Discussion

The p110β and p110δ isoforms of class IA PI3K stand out for their constitutively increased oncogenic and signaling activities. Other isoform-specific properties in class IA include structural features of the p110β–p85β dimer that suggest a unique inhibitory mechanism for the cSH2 domain of p85β [46]. A similar inhibitory activity of the cSH2 domain has also been identified in p110δ [48]. p110β also plays a special role in PTEN negative cancer [24, 49–53]. Expression of the p110δ isoform is restricted to the immune system [54] which is reflected by the involvement of p110δ in hematopoietic malignancies [28, 55]. The congenital mutational activation of p110δ signaling results in the activated PI3Kδ syndrome (APDS), a multifaceted disease that includes immunodeficiency and lymphoproliferation [28, 56, 57]. There have been intense efforts to identify and to test p110δ-specific inhibitors [33, 35, 43, 48, 58–64].

The increased signaling and oncogenic activities of p110β and p110δ have been documented in previous publications [45, 47, 65]. This includes a p110β-specific sequence feature that disrupts the inhibitory interaction between the C2 and iSH2 domains and thus contributes to the elevated activity of p110β [47, 65]. Here we identify a separate, dominant activating sequence motif, PXXP, in the ABD of p110β and p110δ, responsible for the constitutively high activities of both isoforms. Mutating the prolines in the PXXP motif to alanines (P/A mutations) significantly reduces oncogenic and signaling activities of p110β and p110δ. The mutations also weaken the interactions with p85 but do not lead to a complete disruption. The prolines of the PXXP motif are part of the tight ABD–iSH2 binding region. Although they do not contact the iSH2 domain directly, their replacement with alanines would probably result in an interaction that is less tight [46, 60, 66]. Addition of the gain-of-function mutations E552K in p110β and E525K in p110δ to the P/A mutations restores transforming activities but has only a modest positive effect on signaling. These triple mutated proteins (P/A plus helical domain mutations) bind to p85 very weakly, if at all. Single particle cryo-electron microscopy (cryo-EM) analysis of the p110α helical domain mutations E542K and E545K shows a disengagement of p85 domains from p110. Only the iSH2 domain remains tied to the ABD, and the complex of ABD–p85 becomes highly flexible [67]. Data from a hydrogen/deuterium exchange mass spectrometry (HDX-MS) analysis are in accord with the cryo-EM observations but support a less extreme opening of the p110–p85 interface [68]. Biological studies have revealed that deletion of the ABD domain has no significant effect on the activity of the p110α helical domain mutants [69]. In the P/A mutants of p110β and p110δ, the ABD–iSH2 interaction is weakened, and the addition of a helical domain mutation causes a virtually complete separation of p85 from p110. We speculate that these P/A plus helical domain mutations work by the same mechanism as the ΔABD mutants resulting in monomeric p110 with increased expression levels and activity. Previous studies with different cell systems and using different constructs are in accord with the observation that deletion of the ABD leads to increased activity. In human breast epithelial cells, truncation of the ABD in p110α reduces protein stability, but increases activity [26]. In yeast, ABD-deleted p110α and p110β remain enzymatically active [70]. The stability of p110α is dependent on interaction with p85 [71]. However, p110α can show high enzymatic activity without p85 if the stabilizing effect of p85 is mimicked by other conditions [71].

The standard view of the function of p85–p110 interaction is that p85 stabilizes p110 and, in the absence of upstream signaling, inhibits PI3K enzymatic activity [71, 72]. Our data suggest that oncogenic and signaling activities are not critically affected by protein stability and that comparatively unstable p110 or ΔABD p110 constructs can retain high signaling activity.

Could the PXXP sequence interact with proteins that carry an SH3 domain and thus mediate elevated activity? We believe that such an interaction is highly unlikely. The PXXP of p110β and p110δ is not located in a consensus sequence for SH3 binding. Additionally, the PXXP sequence in p110β and p110δ is not linear and therefore cannot accommodate the structure of SH3 domains[73, 74].

The elevated activity of p110β and p110δ can be transferred to p110α by exchanging the ABD of p110α for that of p110β or p110δ. The reciprocal transfer of the alpha ABD to p110β or p110δ reduces activity. Introduction of the PXXP motif in a presumptively analogous site in the alpha ABD enhances activity.

Our data show that the PXXP motif is a dominant determinant of the enhanced activities of p110β and p110δ. In p110β, the PXXP acts probably in concert with the previously identified K342 residue that disrupts inhibitory contacts between the C2 and the iSH2 domains [47]. Mutations of either K342 or PXXP lead to a loss of function. Collectively, the data on p110β and p110δ support the conclusion that the PXXP motif of p110β and p110δ affect the interaction with p85, weakening the inhibitory effect of p85 on PI3K.

## 4. Materials and methods

### 4.1. Cell culture and transfection

Chicken embryo fibroblasts (CEF) were prepared from single, pathogen-free White Leghorn embryos (Charles River, N. Franklin, CT) as previously described [75]. Transfection was performed with avian replication competent retroviral vector RCAS(A) [76] using the polyethylenimine (PEI) transfection method.

### 4.2. Oncogenic transformation

Transformation assays were carried out in six-well plates. CEF were transfected with DNA of the recombinant RCAS(A) constructs using the PEI transfection method and were kept in growth medium for 24 h before being overlaid with nutrient agar containing Ham’s F-10 medium with 20% Earle’s balanced salt solution, 0.6% SeaPlaque Agarose, 3% FBS, 1% heat-inactivated chicken serum, 9% tryptose phosphate broth, 1.8 mM glutamine, 89 U/mL penicillin, 89 μg/mL streptomycin, 1.1% DMSO. This mixture was applied every 2-3 days for 12 days, at which point the overlay was removed and the cell layer stained with crystal violet for quantitative evaluation of oncogenic activity by focus formation [69, 77]. Focus-forming activities depicted in the figures are standardized to 0.5 μg DNA per well.

### 4.3. Constructs and recombinant lentivirus

Vectors with C-terminally FLAG-tagged p110α, p110β, p110δ and their mutants were generated from previously described p110 constructs [45] using the QuikChange XL II site-directed mutagenesis kit (Stratagene, La Jolla, CA). C-terminally FLAG-tagged p110α ABD [32–108], p110β ABD [41–118], p110δ ABD [32–107], p110β ABD [41–118] P/A, p110δ ABD [32–107] P/A, p110α ΔABD [109–1068], p110β ΔABD [119–1070] and p110δ ΔABD [108–1044], p110α [109–1068]-βABD [41–118], p110α [109–1068]-δABD [32–107], p110β [119–1070]-αABD [32–108], p110α [109–1068]-βABD [41–118] P/A, p110δ [108–1044]-αABD [32–108], p110α [109–1068]-δABD [32–107] P/A were generated by PCR, primer sequences are listed in Table 1 (Supplemental information). The mutated genes were subsequently cloned as SfiI DNA fragments into the avian retrovirus vector RCAS(A) [76] and in the lentiviral expression vector pHIV-EGFP (Addgene, Cambridge, MA) with the MCS (HpaI, XbaI, SmaI, and BamHI sites) replaced by modified MCS (SfiI, EcoRI, XbaI and SfiI sites) between the HpaI and BamHI sites. FLAG-tagged p110 and mutants were cloned as SfiI DNA fragments into modified pHIV-EGFP vector. pHIV-EGFP, packing plasmids (Gag, Pol, Rev, and Tat genes) and the envelope protein Env (VSV-G) were transiently co-transfected into 293T cells to generate recombinant virus. All clones were confirmed by protein expression or sequencing.

### 4.4. Western blots and antibodies

CEF were transfected with 0.5 μg of RCAS(A) vector expressing p110α, p110β, p110δ or the specified p110 mutants using the PEI transfection method. After two passages in growth medium, cells were switched for overnight incubation to Ham’s F-10 medium containing 0.5% FCS (calf serum) and 0.1% chicken serum, and this was followed by 2 h in basal F-10. At this point, protein samples were harvested. Western blotting was performed as described [78, 79], with minor modifications. Proteins were extracted from cells using ice-cold RIPA buffer (50 mM Tris·HCl pH 8, 100 mM NaCl, 0.5% Nonidet P-40, 0.5% sodium deoxycholate, 0.1% SDS, 1 mM PMSF, 1 mM NaVO4, and 1× Complete protease inhibitor mixture; Roche, Indianapolis, IN). Proteins were separated by 4-12% gradient SDS/PAGE (Invitrogen, Carlsbad, CA) using the MOPS buffer system. Separated proteins were transferred to PVDF membranes (Millipore, Billerica, MA) with transfer buffer and a transfer apparatus (Invitrogen, Carlsbad, CA). Transferred proteins were visualized with Ponceau S staining, blocked with 5% BSA in TBST. Primary antibodies were added as follows: FLAG (DYKDDDDK)-tag (#14793), HA-tag (#2367), AKT (#4685), AKT p-S473 (#4051), p-S6 ribosomal protein S235/236 (#2211), Total S6 ribosomal protein (#2217), GAPDH (#2118) (Cell Signaling, Danvers, MA). Secondary antibodies were rabbit (No. 31462) or mouse (No. 31432) anti-HRP (Thermo Scientific, Asheville, NC). Western blots were visualized using HRP conjugates, and detection was performed using Super Signal West Pico Chemiluminescent Substrate (Thermo Fisher Scientific, Asheville, NC) according to the manufacturer’s specifications.

### 4.5. Co-immunoprecipitation

293T cells were infected with lentivirus that expresses HA-tagged p85β, p85α, and FLAG-tagged or untagged p110, using an isoform-specific antibody. The p110 isoforms included p110α, p110β, p110δ, p110β P/A, p110δ P/A, p110β P/A with E552K, p110δ P/A with E525K, tagged p110α ABD, p110β ABD, p110δ ABD, p110β ABD P/A, p110δ ABD P/A, p110α ΔABD, p110β ΔABD and p110δ ΔABD, p110α-βABD, p110α-δABD, p110β-αABD, p110α-βABD P/A, p110δ αABD, and p110α-δABD P/A. Lentivirus infections were at a MOI of 1 or higher (confirmed by EGFP expression). Cells were rinsed with phosphate-buffered saline (PBS) and lysed in immunoprecipitation lysis buffer (20 mM Tris-Cl, pH 7.4, 150 mM NaCl, 0.5 mM EDTA, 0.5% NP-40, 1 mM NaVO4, 0.5 mM phenylmethlysulfonyl fluoride (PMSF) and 1× Complete protease inhibitor mixture; Roche, Indianapolis, IN). Cell lysates (100 to 300 μg of protein) were mixed with primary HA-tag antibody (#2367, Cell Signaling, Danvers, MA) and incubated overnight at 4°C with gentle agitation. The lysates were then incubated with protein A/G beads (sc-2003, Santa Cruz Biotechnology, Santa Cruz, CA) for 1 h at 4°C with agitation. The beads were washed three times with washing buffer (20 mM Tris pH 7.5,100 mM NaCl, 0.5% NP-40, 0.5 mM EDTA, 0.5 mM PMSF), bound proteins were eluted by boiling in 4XSDS sample (Invitrogen, Carlsbad, CA) and analyzed for p85-p110 interaction by Western blotting using an antibody against FLAG (DYKDDDDK) tag (#14793) or an isoform-specific antibody. Figures of co-immunoprecipitations depict representative results from at least three repeat experiments.

### 4.6. Protein stability

CEF were transfected with RCAS(A) expressing FLAG-tagged p110δ and p110δ mutants. After two passages, the cells were treated with the translation inhibitor cycloheximide at 50 μg/ml to stop protein translation. Total protein samples were harvested at 0, 2, 4, and 6 h after addition of cycloheximide, and protein levels were determined by immunoblotting using FLAG tag anti- body.

## Supplementary Materials

Table S1: Primer sequences.

## Supporting information

Table S1: Primer sequences

## Acknowledgment

The authors want to express their gratitude to Marc Elsliger for illuminating discussions on the p110 - p85 interaction.

## Author Contributions

Conceptualization, P.K.V. and S.K.; methodology, Y.I.; validation, P.K.V., J.R.H. and Y.I.; formal analysis, P.K.V. and J.R.H.; investigation, Y.I., P.P., C.S.L., L.U.; resources, S.K., Y.I., L.U.; data curation, J.R.H.; writing – original draft preparation, Y.I., J.R.H. and P.K.V.; writing – review & editing, P.K.V.; visualization, P.K.V., J.R.H. and Y.I.; supervision, P.K.V.; project administration, P.K.V.; funding acquisition, P.K.V. All authors have read and agreed to the published version of the manuscript.

## Funding

This work was supported by the National Institutes of Health under awards R35 CA197582, T32 DK007022 (PP) and T32 AI007354 (CSL). The content is solely the responsibility of the authors and does not necessarily represent the official views of the National Institutes of Health.

## Conflicts of Interest

The authors declare no conflicts of interest.

## Notes

### Competing Interest Statement

The authors have declared no competing interest.

