## Supplementary material for "The constitutive oncogenic and signaling activities of phosphatidylinositol 3-kinase (PI3K) isoforms p110β and p110δ": Table S1: Primer sequences

```
TABLE 1

P110
Alpha
(Forward) 	AtatGGCCATTACGGCCATGgaagaaaagatcctcaatcgagaaattgg
(reverse) 	TCActtatcgtcgtcatccttgtaGTTCAATGCATGCTGTTTAATTGTGTGGAA

Beta
(Forward) 	AtatGGCCATTACGGCCATGggggaaaaattagactcaaaaattggagtc
(Reverse) 	TCActtatcgtcgtcatccttgtaAGATCTGTAGTCTTTCCGAACTGTGTGGGC


Delta
(Forward)   	  AtatGGCCATTACGGCCATGgtgaagaagctcatcaactcacagatca
(Reverse) 	TCActtatcgtcgtcatccttgtaCTGCCTGTTGTCTTTGGACACGTTGTGGGC


P110 w/o ABD
ALpha
(Forward)	 	AtatGGCCATTACGGCCATGgaagaaaagatcctcaatcgagaaattgg

Beta
(Forward) 	AtatGGCCATTACGGCCATG ggggaaaaattagactcaaaaattggagtc

Delta
(Forward) 	AtatGGCCATTACGGCCATG gtgaagaagctcatcaactcacagatca
```

```
P110 ABD SWAPPED
P110 DELTA ABD / P110 ALPHA
(FORWARD)		taaaagtaattgaaccagtaggcaaccgtcgcgtgaagaagctcatcaactcacagat
(REVERSE) 	ATCTGTGAGTTGATGAGCTTCTTCACGCGACGGTTGCCTACTGGTTCAATTACTTTTA

P110 ALHA ABD / P110 DELTA
(FORWARD) 	tcctgcgcctggtggcccgtgagggcgacgaagaaaagatcctcaatcgagaaattggt
(REVERSE)		ACCAATTTCTCGATTGAGGATCTTTTCTTCGTCGCCCTCACGGGCCACCAGGCGCAGGA

P110 BETA ABD / P110 ALPHA
(FORWARD) 	taaaagtaattgaaccagtaggcaaccgtggggaaaaattagactcaaaaattggagtc
(REVERSE)		GactccaatttttgagtctaatttttccccACGGTTGCCTACTGGTTCAATTACTTTTA

P110 ALPAH ABD / P110 BETA
(FORWARD)		ctcaaattagtgacaagaagttgtgacccagaagaaaagatcctcaatcgagaaattggt
(REVERSE) 	ACCAATTTCTCGATTGAGGATCTTTTCTTCtgggtcacaacttcttgtcactaatttgag

P110 abd DOMAIN
(REVERSE)		TCActtatcgtcgtcatccttgtaACCAATTTCTCGATTGAGGATCTTTTCTTC

(REVERSE)		TCActtatcgtcgtcatccttgtaGactccaatttttgagtctaatttttcccc

(REVERSE)		TCActtatcgtcgtcatccttgtaATCTGTGAGTTGATGAGCTTCTTCACGCG
```
